# The structure of species discrimination signals across a primate radiation

**DOI:** 10.1101/574558

**Authors:** Sandra Winters, William L. Allen, James P. Higham

## Abstract

Discriminating between conspecifics and heterospecifics potentially challenging for closely related sympatric species. The guenons, a recent primate radiation, exhibit high degrees of sympatry and form multi-species groups in which hybridization is possible but rare in most populations. Guenons have species-specific colorful face patterns hypothesized to function in species discrimination. Here, we apply a novel machine learning approach to identify the face regions most essential for correct species classification across fifteen guenon species. We then demonstrate the validity of these computational results using experiments with live guenons, showing that facial traits identified as critical for accurate classification do indeed influence selective attention toward con- and heterospecific faces. Our results suggest variability among guenon species in reliance on single-trait-based versus holistic facial characteristics when discriminating between species, and differences in behavioral responses to faces can be linked to whether discrimination is based on a single trait or whole-face pattern. Our study supports the hypothesis that guenon face patterns function to promote species discrimination and provides novel insights into the relationship between species interactions and phenotypic diversity.

## Introduction

Closely related species living in sympatry face a potential challenge in discriminating between conspecifics and heterospecifics. Such decision-making has important selective outcomes, particularly in behaviors such as mate choice, with individuals choosing heterospecific mates often incurring substantial fitness costs [1]. One mechanism for avoiding the costs if interacting with heterospecifics is the use of species-specific signals that structure behavioral interactions between species. For instance, mating signals and associated mating preferences that differ between sympatric heterospecifics can function to maintain reproductive isolation across species boundaries [2]. Such signals are predicted to be salient and distinctive [3], with sympatric species under selective pressure to diversify. A pattern in which signal distinctiveness increases with degree of sympatry, known as character displacement [4,5], has been observed in a wide variety of animal groups [6–13]. Importantly, signals that function to maintain reproductive isolation via mate choice should elicit increased mating interest from conspecifics compared to heterospecifics [14].

Species in evolutionarily young animal radiations may be at particular risk of hybridization and other costly interactions with heterospecifics due to behavioral similarities and a lack of post-mating barriers to reproduction [15]. One such radiation is the guenons (tribe Cercopithecini), a group of African primates consisting of 25-38 recognized species [16–18] that diverged from papionin primates around 11.5 million years ago [19]. Guenons exhibit high degrees of sympatry and often form polyspecific groups in which multiple species travel and forage together [20]. Many guenons therefore interact with heterospecifics that share general patterns of morphology (e.g. overall body size/shape) and behavior (e.g. activity patterns). In such circumstances, discriminating between con- and heterospecifics may be particularly important, especially in a mating context. Hybridization between sympatric guenon species is possible but rare in natural circumstances [21], suggesting the existence of barriers to heterospecific mating within mixed-species groups.

Guenons are among the most colorful and visually patterned groups of primates with many species exhibiting extraordinary and unique face markings [10,23,25–27], which are minimally variable between sexes across all guenon species [23,24]. Kingdon [23,26,27] hypothesized that guenons use their divergent facial appearances to distinguish between species and therefore select appropriate mates. This young and impressively diverse primate radiation represents a fascinating test case of how visual signals are involved in species radiations and mixed-species interactions [5,28–30]. Recent empirical work has begun to generate evidence for their key role in guenon phenotypic and species diversification. Images of guenon faces can be reliably classified by species using computer algorithms [10,24], demonstrating that guenon faces contain species-specific identifying information. Guenon face patterns also exhibit character displacement, with facial distinctiveness between species increasing with degree of sympatry across the group [10]. Moreover, facial components common across species (nose spots and eyebrow patches) alone can be used to computationally classify species [24]. This suggests that guenon faces may be somewhat modular, with species information encoded in particular face regions. Which face regions are most important, and the extent to which such regions vary across species remains an open question that is of key importance to understanding how complex signals involved in species discrimination evolve. Critically, it is unknown whether variation across guenon species in purported species discrimination signals is perceived and acted on by con- and heterospecific receivers.

Here, we use a machine learning approach to identify guenon face regions that are most important for correct species classification by a computer. These results objectively identify the signal components most likely to be useful to guenon receivers. We use them to determine which signal properties to systematically investigate in behavioral experiments with guenon observers. The machine-learning stage is critical, as many experiments that investigate behavioral responses to complex signals select manipulations based on the perceptions of investigators, which introduces anthropocentric bias [31]. Using the guenon face image database produced by Allen et al. [10], we couple eigenface decomposition of the faces [32] with a novel occlude-reclassify scheme in which we systematically block each part of the face and reclassify the image. This allows us to document the spatial distribution of species-typical information across guenon faces by identifying which face regions, when obscured, cause the break-down of correct species classification. Eigenface decomposition was originally developed for individual face discrimination in humans [32]; feature detection based on eigenfaces is also applicable to other types of discrimination tasks involving complex animal signals [33–35] and has been used previously to quantify guenon facial variation [10]. The perceptual face space generated by eigenface decomposition parallels mammalian visual processing [36], lending biological credibility.

After identifying the face regions that cause break-down in classification, and thus those that should be important for correct species identification, we then present captive putty nosed monkeys (*Cercopithecus nictitans*) and mona monkeys (*C. mona*) with images of con- and heterospecific faces exhibiting variation in these regions and measure their resulting eye gaze to assess their ability to distinguish between species based on face patterns. Ours is the first direct measure of guenon responses to con- and heterospecific faces, which is crucial for clarifying the biological relevance of guenon face patterns and for validating previous correlational results. Differences in looking time between classes of stimuli can be difficult to interpret due to various and often unpredictable novelty and familiarity effects [37], however primates reliably exhibit a visual bias (i.e. greater looking time) toward images of conspecifics compared to those of heterospecifics [38–42]. We follow the interpretation that longer looking time at a particular face reflects level of interest. This is consistent with an interpretation that the face resembles a conspecific face more closely, though other explanations are possible.

Our experimental trials involve the simultaneous presentation of paired con- and heterospecific faces, focusing on a particular facial trait for each species. For putty nosed monkeys we focus on nose spots and for mona monkeys on eyebrow patches, on the basis that each of these features is within the region of the face identified by our machine learning approach as being critical for that species. In each trial, heterospecific faces either do or do not share a focal face trait with the subject, and conspecific faces are presented either naturally or after being modified to remove the focal trait (for example stimuli, see Figure 1). This approach allows us to assess generalized species biases in degree of interest as well as the extent to which particular face regions influence these biases.

**Figure 1.**
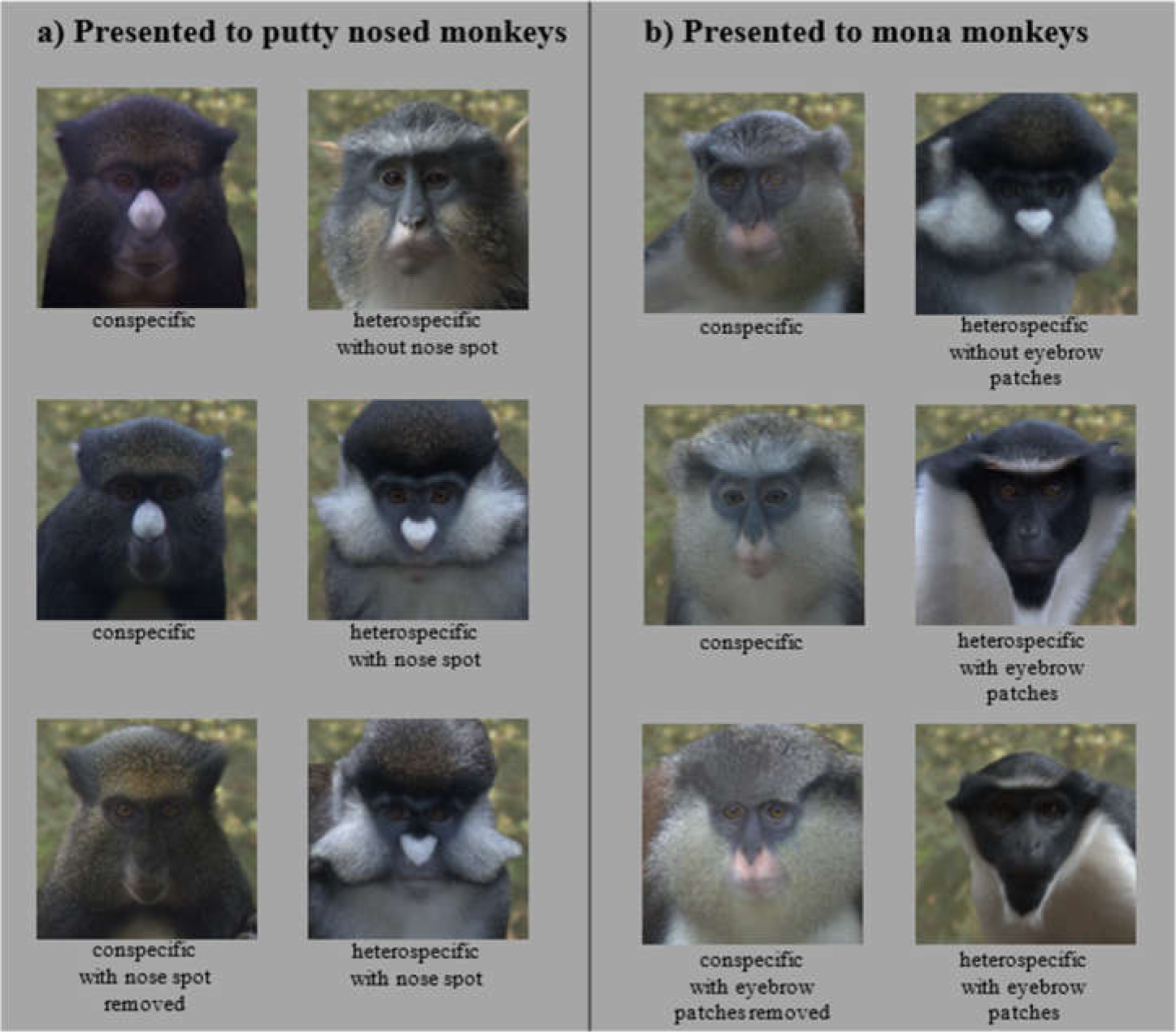
Example experimental stimulus pairs. Subjects were shown a pair of stimulus images consisting of a conspecific and a heterospecific. Facial traits (nose spots for putty nosed monkeys and eyebrow patches for mona monkeys) were varied across trials, with conspecifics paired with a heterospecific species that shares the facial trait (row 1) and one that does not (rows 2 and 3). Conspecifics were displayed either naturally (rows 1 and 2) or with the facial trait removed (row 3). All subjects participated in all three trial types. Trial order and stimulus image side were counterbalanced across subjects.

We predicted variability across species in the face regions identified by our occlude-reclassify procedure, but made no predictions regarding which regions in particular would be essential for each species. In looking time experiments, we predicted that putty nosed and mona monkeys would exhibit visual biases toward face images of conspecifics, and that these biases would be influenced by species-typical facial characteristics identified as important for correct species classification. Such a pattern of results would support a role for species discrimination signals likely used to facilitate inter-specific interactions such as maintaining reproductive isolation via mate choice in generating and maintaining phenotypic variation in one of the most speciose and diverse primate radiations. Species discrimination requires differentiating between conspecifics and all heterospecifics at the very least, however it may be possible that animals may also be able to discriminate between different heterospecific species. Ultimately, by examining how aspects of highly complex signals encode species identity and influence receiver biases, this research increases our understanding of how selection for species identity signaling generates phenotypic diversity.

## Methods

### Image collection & processing

Guenon face pattern analyses are based on an existing database of guenon face images from 22 guenon species [10]. Detailed methods of image collection and processing have been published elsewhere [10]. Briefly, we used digital images of captive guenons collected using a color-calibrated camera. Multiple images were taken of each subject while in a front-facing position under indirect light. Images were transformed from camera RGB color space to guenon LMS color space, defined by the peak spectral sensitivities of guenon long, medium, and short wavelength photoreceptors. All images were then standardized with respect to illumination, size, blur, and background. Each image was resized to be 392 by 297 by 3 pixels. All pixel values were represented using double-level precision.

To avoid classifying species based on a very small number of exemplars, we restricted our analyses to species represented by at least four individuals in our image database (i.e. all classifications in a leave-one-out procedure are made based on at least three exemplars; see below). Our analysis is therefore based on 599 total images of 133 individuals, collectively representing fifteen guenon species (for species-specific sample sizes, see Figure 3).

### Identification of face regions important for species classification

Guenon face images can be reliably classified by species based on eigenface features [10,24]. This approach relies on dimensionality reduction via principal component analysis (PCA) to extract relevant features from face images; these features can then be used for the classification of new faces [32]. In this procedure, each ‘eigenface’ (i.e. the eigenvectors resulting from PCA of all face images) represents a different dimension of facial variability and each face image can be represented by a series of weights associated with each eigenface. This creates a multi-dimensional ‘face space’ in which faces are represented as points based on their eigenface weights, and zero weights for all eigenfaces (i.e. the center of the space) represents the average face across all images. Such face spaces have psychophysical parallels in primate face processing centers in the visual cortex [36]. Multiple images of each subject were averaged to generate average individual faces, which in turn were used to generate the average species faces that were used in eigenface decomposition. We classified new images using a nearest-neighbor classifier based on minimum Euclidean distance to each average species face in face space. This scheme corresponds to an average face model of guenon face learning, which assumes that guenons cognitively encode different species’ face patterns as the mean of all encountered examples. In previous work using similar methods, results were robust to the choice of learning model [10].

To avoid using the same individual guenons to both train and test our species classifier we used a leave-one-out procedure for all analyses. For this procedure, we systematically removed each individual from the image set, repeated the analysis procedure outlined above, then classified each image of the excluded individual based on the features generated from all other images. All species included in these analyses are represented by at least four individuals (range: 4-23). We present results for all species, however results for species with samples sizes in the lower end of this range should be considered less robust and interpreted with caution.

Eigenface-based features can be used to reliably classify guenons by species based on axes of variation, however the extent to which specific facial characteristics are relevant for correct classification of each species is difficult to determine. We used an occlude-reclassify scheme developed to identify which image regions contribute most to correct classification in computer vision classification tasks [43]. For each correctly classified image, we systematically blocked each image region and re-classified the image; a correct re-classification indicates that the occluded region of the face was unnecessary for correct classification, while an incorrect re-classification indicates that the occluded region was essential. Occlusion of face regions was accomplished by setting the relevant pixel as well as all those in a thirty-pixel radius to the mean face color of that species. This procedure was repeated for every pixel in the image, effectively sliding the occluded region across all face areas. A radius of thirty pixels occludes approximately five percent of the image (Figure 2), with the specific region being occluded shifting by one pixel at each iteration. Primate faces are broadly symmetrical, therefore to avoid the presence of duplicate spatial information that may support species classification when part of the face is occluded, we ran analyses on the left and right halves of the face separately. Results differed little, so for clarity we report the results from the left hemi-face classification in the main text, with right-side results summarized in the supplementary results. For more details on the implementation of the occlude-reclassify procedure, see supplementary methods. Based on this occlude-reclassify scheme, we generated a binary image for each image in our data set, with each pixel being either zero (black) or one (white) base on whether the image was correctly classified when that pixel and its neighbors was occluded. We then averaged these binary images across individuals and species to generate species level heatmaps depicting face regions that are essential for correct classification across species. For visualization, we converted greyscale heatmaps to color using a color mapping function. To facilitate the identification of critical face regions, occlusion results are presented as composite images combining heatmaps and a greyscale version of the relevant species average face, with transparency set to 0.5.

**Figure 2.**
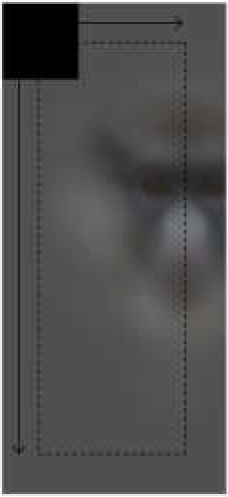
Average guenon face with an occluder shown in the top left. The occluder is depicted in black for maximal visibility, but in analyses presented here is set to the mean face color of the relevant species. During the occlude-reclassify analysis, the occluder is slid across the image and the image re-classified; an incorrect classification at a given occluder location indicates the presence of face information critical to correct classification. Analyses are run on hemi-faces to account for facial symmetry. Image borders outside the radius of the occluder are not tested; the dashed line encloses the region of the image analyzed using the occlude-reclassify procedure.

Heatmaps vary across species in the extent to which face regions identified as essential for correct species classification are spread across the face (i.e. ranging from small and isolated face regions to large portions of the face identified as critical) as well as the relative import of identified regions (i.e. the likelihood that identified regions caused misclassification, encoded as how dark identified regions are in the heatmap). To quantify the spread and relative importance of the identified face regions across species, we calculated the proportion of the face misclassified and the mean classification error, respectively. The proportion of the face misclassified was calculated as the number of heatmap pixels less than one (i.e. those that were ever incorrectly classified) divided by the total number of pixels in the average face for each species; higher values indicate that the face regions essential for correct species classification are spread more widely across the face. The mean classification error was calculated as the mean value of all heatmap pixels less than one; higher values indicate that the face regions identified are particularly critical and more often lead to misclassification when occluded (i.e. the identified regions are darker in the heatmaps). Computational analyses were conducted in MATLAB™ and run on the High Performance Computing cluster at New York University.

### Looking time experiments

Looking time experiments were conducted at CERCOPAN sanctuary in Calabar, Nigeria, and included 18 adult putty nosed monkeys (6 males, 12 females) and 16 adult mona monkeys (10 males, 6 females). Each species was divided into four experimental groups (based on socially housed groups), with all individuals in the group viewing the same images in the same order. In each species, two experimental groups were presented with male stimulus images and two with female stimulus images across all trials. Experiments involved the simultaneous presentation of two stimulus images to subjects, with their resulting eye gaze measured to determine visual biases. Stimulus preparation and experimental procedures were carried out following the recommendations of Winters et al. [37]. Briefly, we prepared stimulus images depicting guenon faces which were presented approximately life-sized (image size on screen: 500 × 500 pixels, 11.96 × 11.96 cm), with accurate colors, and standardized for relevant characteristics. Stimulus image pairs were presented to subjects side-by-side using a custom-designed experimental apparatus. For more details regarding subjects, stimuli preparation, and experimental apparatus design, see supplemental methods.

Each subject participated in three trials, with stimulus image pairs depicting the following: (1) a conspecific and a heterospecific that shares a focal trait with the conspecific, (2) a conspecific and a heterospecific that does not share a focal trait with the conspecific, and (3) a conspecific for which the focal trait has been modified and a heterospecific that shares the focal trait with the conspecific. Heterospecifics presented to putty nosed monkeys were Wolf’s guenons (*C. wolfi*, no nose spot) and red-tailed monkeys (*C. ascanius*, nose spot); heterospecifics presented to mona monkeys were red-tailed monkeys (no eyebrow patches) and Diana monkeys (*C. diana*, eyebrow patches). Heterospecific species were selected based on the presence/absence of the relevant facial trait, a lack of range overlap with the subject species, and availability of sufficient and appropriate images in our database. Image presentation locations (i.e. left verses right) were counterbalanced across trials, and trial order was varied across subjects; both factors were included in statistical analyses. For each trial, we placed the experimental apparatus immediately outside the relevant enclosure and recorded the identities of participating subjects. We waited a minimum of one week between trials of the same subject to minimize habituation or trial order effects.

Videos of each trial were coded frame by frame to quantify the amount of time subjects spent looking at each stimulus image. All coding was done blind to trial conditions and stimulus image location. Reliability was assessed using approximately 10% of all trial videos, in which we assessed agreement between two coders on the direction of jointly coded looks within these trials as being in agreement in 94.46% of frames (Cohen’s kappa = 0.883), which is well within the range of acceptable reliability scores for this type of data [37,44]. Raw looking time data was compiled to yield a total number of frames spent looking at each stimulus image for each subject in each trial. Subjects varied widely in their level of interest in experiments, resulting in considerable variation in overall looking time. We therefore used only the first five seconds of looking for each subject in each trial, while allowing them to complete the current look at the five second mark (i.e. we required at least one second of non-looking before terminating coding for each subject). This resulted in a mean total looking time (± standard deviation) of 3.89s (± 1.98s) for putty nosed monkeys and 4.58s (± 2.52s) for mona monkeys, which is similar to durations reported in previous looking time experiments in primates [37,44]. Because a direct comparison is made between the species depicted in stimuli, each trial effectively serves as its own control.

### Statistical analyses

We analyzed differences in looking time elicited by subjects in experimental trials using generalized linear mixed models (GLMMs). Models were fit using a binomial family distribution, with the number of video frames spent looking at the targeted stimulus image and the number of video frames spent looking at the paired image set as the binomial outcome variable. This structure allowed us to assess looking biases while accounting for any differences in total looking time across subjects. All models included group, subject, and unique trial (i.e. a unique identifier for each subject in each trial, included to account for our analysis of the two images presented in each trial as separate data ‘rows’) as nested random effects. Stimulus species (conspecific v. heterospecific) and focal trait similarity (presence of nose spots for putty nosed monkeys and eyebrow patches for mona monkeys), were included as fixed effects. We also included the following additional factors as fixed effects: subject age (log transformed), sex, and origin (captive v. wild born); stimulus image presentation spot (right v. left), eye contact (direct eye contact with the camera or looking slightly away), sex, and degree of familiarity to the subject; and trial order, apparatus pattern, and display ICC profile. For more details about these variables see supplemental methods.

To determine which variables significantly influenced subject looking biases, we compared models with different parameterizations using likelihood ratio tests (LRTs). A single model including all fixed effects simultaneously would involve an excessive number of predictors. We therefore first analyzed each variable separately via comparisons to a null model including only random effects, and excluded non-significant predictors from subsequent analyses. We generated an initial model composed of factors that were statistically significant (alpha < 0.05) or exhibited a trend (alpha < 0.1) when tested alone. To determine the statistical significance of these factors we then systematically excluded each factor from this model and tested its contribution to the fit of the model to the data using LRTs. When species (conspecific v. heterospecific) and focal trait (shared v. not shared) were both significant predictors in this model we also tested a species*trait interaction. Within a final model composed of significant predictors we compared across factor levels of fixed effects using z scores calculated using a normal approximation. Adherence to model assumptions was verified based on plots of fitted values and residuals. Trials from putty nosed and mona monkeys were analyzed separately. GLMMs were run using the ‘lme4’ package version 1.0.12 [45] in R version 3.3.3 [46].

## Results

### Occlude-reclassify machine classification

We began by confirming that guenons could be reliably classified by species based on eigenface decomposition [10]. Average subject images were correctly classified by species 99.31% of the time, and distinct images were correctly classified 93.03% of the time. All correctly classified images (n = 654) were used to identify face regions of critical importance to correct species classification by the computer algorithm, using our occlude-reclassify scheme. We identified essential face regions in all guenon species that, when occluded, led to incorrect species classification (Figure 3; for full resolution images see Supplementary File 1). Species differed in the importance of different face regions as well as the extent to which important regions were concentrated in specific facial features or were more widely distributed across larger face areas (Figure 4). For example, the nose spot of the putty nosed monkey was the most critical facial feature identified across all species. The putty nosed monkey had the highest mean error rate for misclassified face regions – indicating that the face regions identified had the highest likelihood of causing misclassification when occluded – with the essential regions centered exclusively on the nose. Thus, in the putty nosed monkey the nose is the only essential face feature; when the nose is occluded species classification breaks down, whereas occluding any other face region has no effect. In contrast, in other species our classifier relied on broader regions of the face, with larger face regions identified as important for correct classification and the classifier relying less exclusively on a single feature. The mona monkey is a good example of this, with disparate face regions including the cheeks, eyebrows, and ear tufts all influencing correct classification of this species. In some species negative space is important, suggesting that what makes the faces of species distinctive may be the absence of certain facial traits. For instance, in *M. talapoin* the absence of distinctive traits along the sides of the face – such as cheek and/or ear tufts observed in other species – appears to be important.

**Figure 3.**
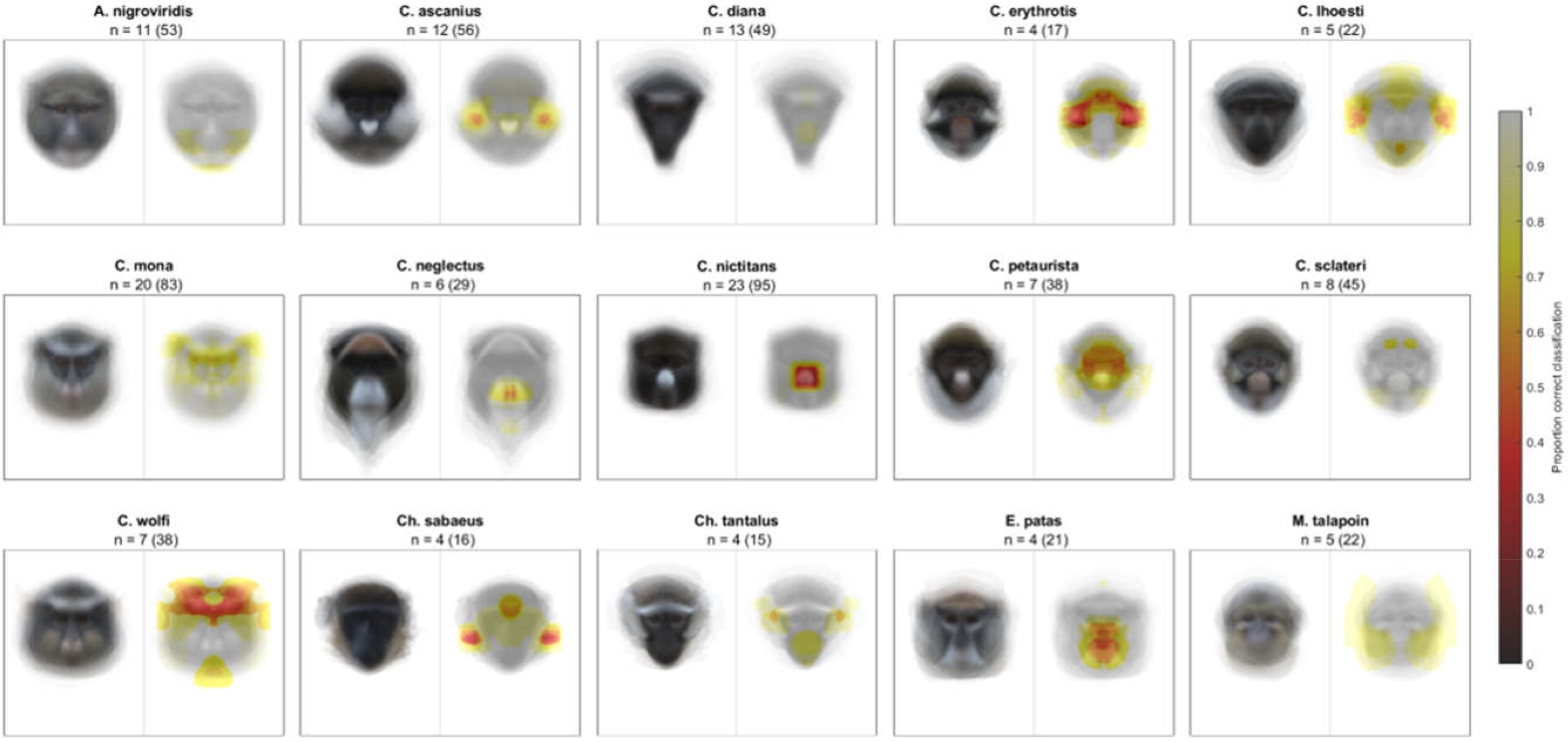
Likelihood of correct classification based on occlusion of different face regions. Species average faces are displayed on the left and heatmaps identifying critical face regions on the right. Sample size is reported as n = number of individuals (number of total images).

**Figure 4.**
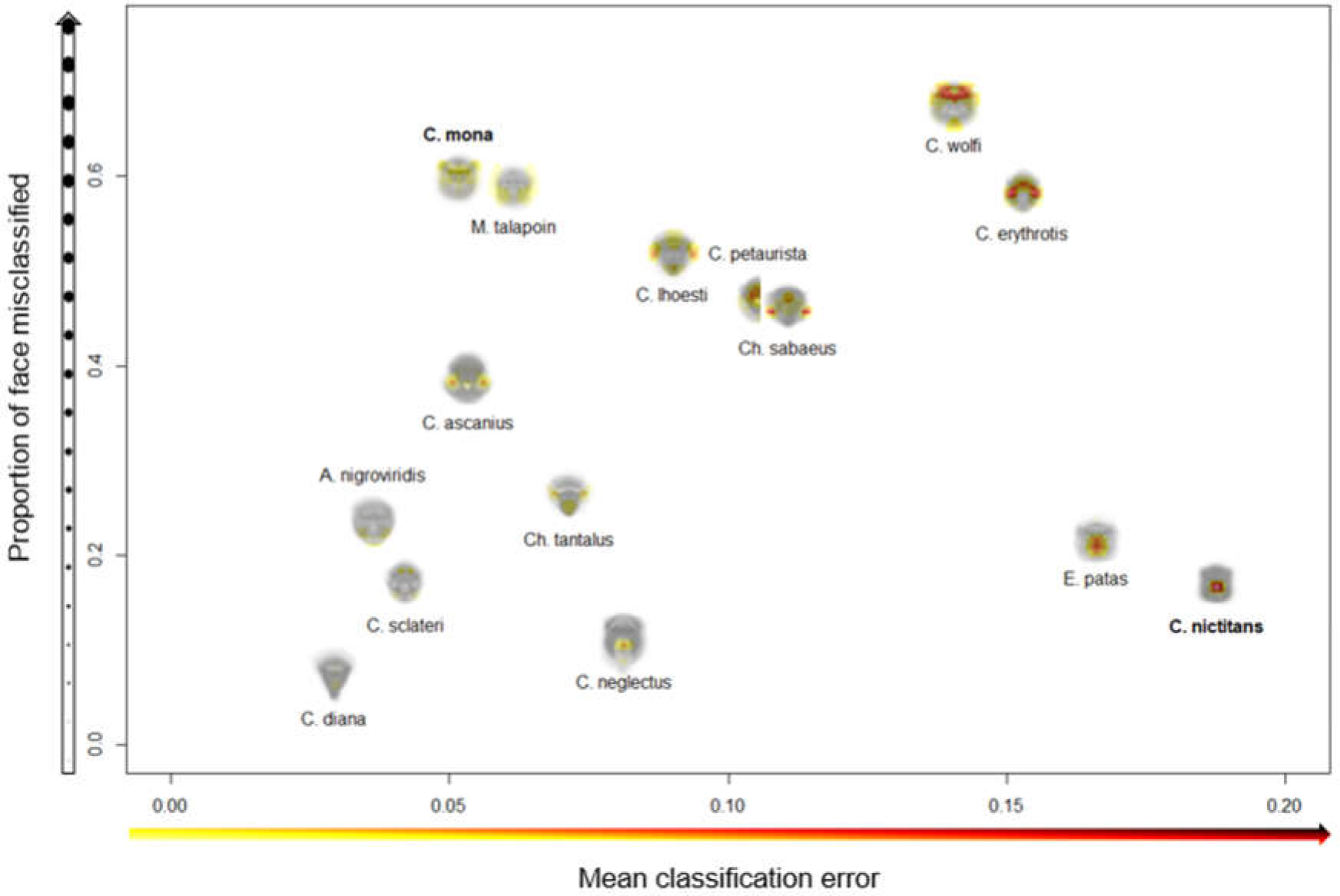
Variation across species in face regions identified as essential for correct species classification. The proportion of the face misclassified (y-axis) indicates the spread of essential regions across the face; higher values signify broader spread and lower values more concentrated regions. The mean classification error (x-axis) measures the relative importance of identified features; higher values indicate higher rates of misclassification, suggesting identified regions are particularly essential for correct species classification. Experimental results are presented for C. mona and C. nictitans (Figure 5).

### Looking time experiments

Our experiments presenting subjects with pairs of con- and heterospecific faces revealed visual biases in resulting eye gaze in both putty nosed and mona monkeys. In the subset of trials that included a natural conspecific and a heterospecific without the relevant face trait (i.e. those where the relevant facial traits are not spread across both con- and heterospecific faces), species (and therefore also facial trait) was a significant predictor of looking behavior (putty nosed monkeys: Chisq = 63.312, p < 0.001; mona monkeys: Chisq = 30.755, p < 0.001), with both putty nosed and mona monkeys exhibiting a conspecific bias (respectively: z = 7.920, p < 0.001; z = 5.536, p < 0.001; Figure 5).

**Figure 5.**
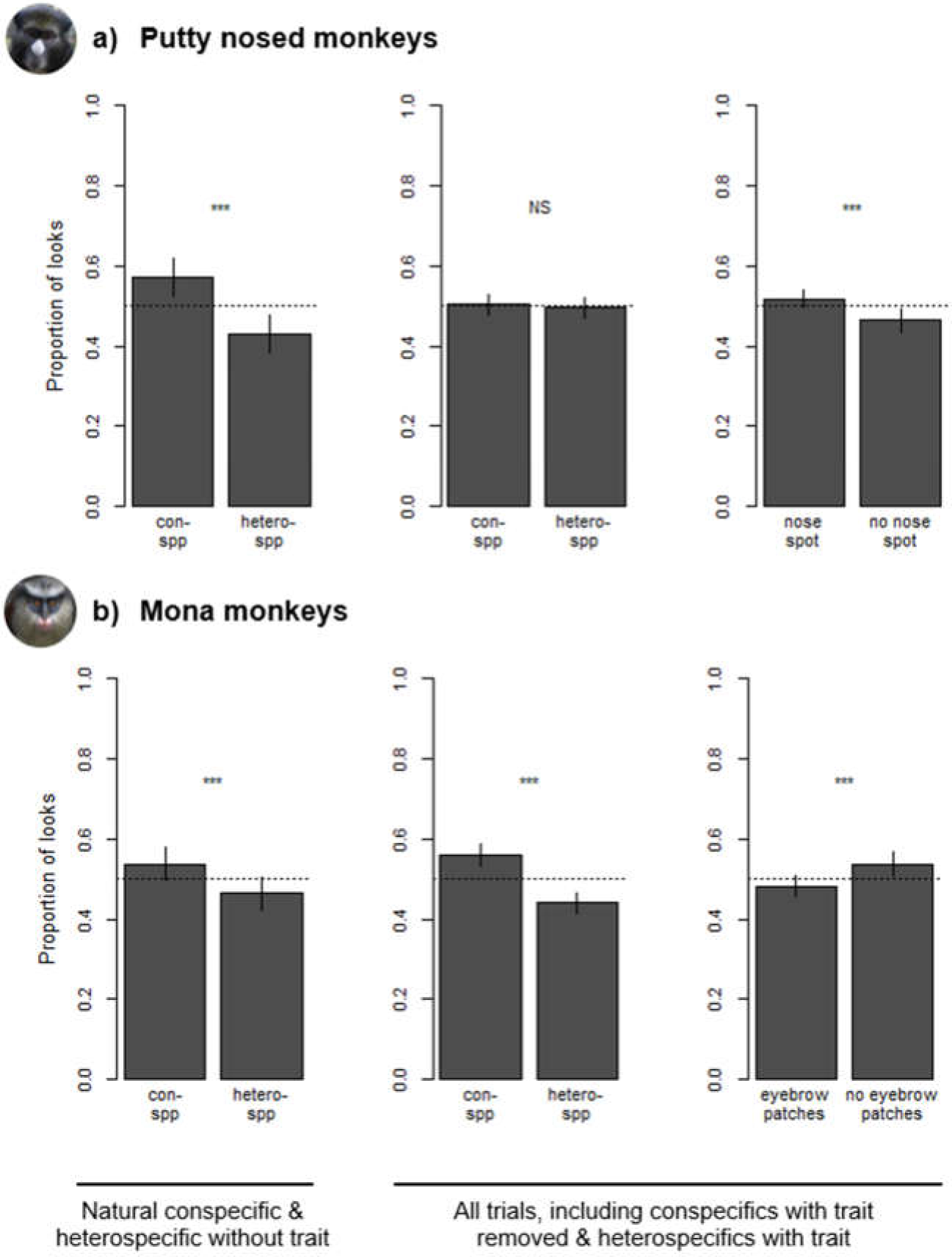
Species and trait biases observed during looking time tasks with (a) putty nosed monkeys and (b) mona monkeys. Leftmost plots depict differences in looking time in trials consisting of conspecifics and heterospecifics without the relevant facial trait. Center and right plots depict looking time differences across all trials – which also include heterospecifics with the relevant facial trait and conspecifics without it – with species biases depicted in the center and trait biases on the right. Results are based on 18 putty nosed monkeys and 16 mona monkeys. Each subject participated in three trials (see Figure 1 for example stimuli for each trial type). Error bars indicate the standard error of the mean.

Across all trials, in putty nosed monkeys model comparisons revealed that looking behavior was significantly influenced by facial trait (nose spot v. no nose spot; Chisq = 11.511, p < 0.001) and image location (right v. left; Chisq = 18.065, p < 0.001), but not by species (conspecific v. heterospecific; Chisq = 3.051, p = 0.081). Overall, putty nosed monkeys looked longer at stimulus faces that displayed a white nose patch (z = 3.343, p < 0.001; Figure 5), their diagnostic species trait, regardless of species identity. Putty nosed monkeys also exhibited a significant right gaze bias (z = 4.289, p < 0.001). None of the other variables relating to subject, stimulus, or trial characteristics were statistically significant (all p > 0.1; Supplementary Table 1).

In mona monkeys, model comparisons revealed that looking behavior was significantly influenced by species (conspecific v. heterospecific; Chisq = 177.480, p < 0.001), facial trait (eyebrow patches v. no eyebrow patches; Chisq = 29.462, p < 0.001) and a species*trait interaction (Chisq = 8.242, p = 0.004). Across all trials, mona monkeys looked longer at conspecifics (z = 9.945, p < 0.001; Figure 5) and as a separate effect, faces without white eyebrow patches, one component of their overall wider diagnostic discrimination area (z = 5.851, p < 0.001). There was also an interaction between these two variables, with mona monkeys looking longer at heterospecific faces with white eyebrow patches (z = 2.868, p = 0.004). None of the other variables relating to subject, stimulus, or trial characteristics played a significant role in mona monkey visual biases (all p > 0.1; Supplementary Table 2).

## Discussion

Our experiments show that eye gaze in guenons is influenced by face regions identified as critical to correct species classification by our machine classifier. This convergence of results using disparate methods reinforces the validity of both, and ties computationally derived results directly to guenon perception, demonstrating the utility of machine learning for identifying biologically relevant signal components. To our knowledge, ours is the first analysis to use machine classification combined with the systematic occlusion of image regions to characterize the relevant signaling information encoded in an animal’s appearance. This approach, based on research in the field of computer vision designed to assess the contribution of image contents to object classification [43], is useful for objectively quantifying the relative roles of different signal components with respect to overall signal function. In closely-related sympatric species, selection against mating or interacting with heterospecifics is often associated with the evolution of species-typical traits used to maintain reproductive and behavioral isolation. The guenons, a recent and diverse radiation that exhibit mixed species groups in which hybridization is rarely observed, exemplify this phenomenon. By showing how species classification is dependent on different aspects of face patterning and that this links with looking time toward con and heterospecifics, our analyses support a role for guenon face patterns in species discrimination, and identify specific face regions critical for this function. This parsing of critical signal components is critical for understanding the phenotypic evolution of complex signals and identifying relevant axes of signal variation for additional analyses.

Our occlude-reclassify analysis identified face regions critical to correct species classification by a machine classifier in all guenon species included in our study. Critical regions differed in both location and spread across the face, suggesting variation in potential use across species. For some guenons, reliance on a single facial characteristic may be sufficient for species discrimination. The best example of this in our data set is the putty nosed monkey, where our machine classifier relied exclusively on the white nose spot to classify this species. That is, occlusion of any other region resulted in correct classification, but when the nose spot was occluded classification failed. This result is reinforced by our experiments, in which putty nosed monkey visual attention was driven wholly by the presence of nose spots. Putty nosed monkeys exhibited a conspecific bias when presented with natural con- and heterospecific faces, as is typical in primates, however including stimuli depicting heterospecifics with nose spots and conspecifics without nose spots completely obscured this conspecific bias. This combination of results illustrates the importance of nose spots in this species. It is worth noting that putty nosed monkey nose spots are the most straightforward facial trait documented in our analysis (i.e. putty nosed monkeys were only misclassified when the nose spot was occluded and occluding the nose spot led to a high rate of misclassification) and the relative simplicity of the face and related visual biases in this species is likely exceptional. On the whole, species discrimination signals in a large radiation with varying patterns of sympatry are expected to be complex and multidimensional, and it is likely that only some species can exhibit single-trait-based signals and visual biases without the system breaking down. This is supported by our results showing that for most guenon species our classifier relied on multiple face regions for species discrimination.

Not all guenons exhibited critical face regions restricted to a single facial trait, and our machine classifier sometimes relied on disparate face regions. In our data set, the mona monkey is a good example of such a species. Like in putty nosed monkeys, our experiments with mona monkeys supported these computational results. Mona monkeys exhibited a conspecific bias across all trials, regardless of single trait manipulations, as well as an additional bias based on the presence of eyebrow patches. Thus, eyebrow patches alone do not appear to be the sole focus of attention in mona monkeys. We predict that additional manipulation of other face regions would be necessary to redirect their visual attention. Nonetheless, that mona monkey attention is still influenced by this species-typical trait shows that it is important but not essential, a result predicted by our computational analyses. It is unclear why mona monkeys would look longer at stimuli without eyebrow patches, however it is possible that utilization of the whole face causes increased attention to incongruency (e.g. conspecifics without eyebrow patches or heterospecifics with them). Our results suggest that in mona monkeys, species discrimination may be based on broader face information, and the perceptual processes involved in assessing potential mates could be similar to generalized holistic face processing mechanisms observed in other primates [47].

Our results suggest that guenons, while united by a general pattern of facial diversification and the probable use of faces in mate choice, may vary across species in the specific traits and processes that are involved in discriminating between conspecifics and heterospecifics. Our pattern of results for putty nosed monkey nose spots and mona monkey eyebrow patches is interesting because we know that both traits do contain sufficient information to discriminate between species that share these features [24], yet they influence attention differently in the two species. This disparity highlights the importance of testing receiver perception directly. The fact that our experimental results with guenons line up with predictions generated by our occlude-reclassify analysis implies that these computationally derived results are biologically valid. Interestingly, we found no sex differences in visual biases for either species, suggesting that selective pressures on species discrimination signaling and preference traits are similar between sexes.

In guenons, an observed lack of hybrids in most polyspecific groups [21] is notable given that hybridization is known to be possible between many guenon species [21,23,27], and indicates the existence of pre-mating barriers to reproduction. Increased eye gaze is associated with increased mating interest in humans [48] and non-human primates [44,49], suggesting that our experimental results would generalize to mating contexts in guenons. Combined with previous work [10,23,24,26,27], our results support the hypothesis that guenon face patterns play a role in mate choice and reproductive isolation in this group. However, it remains possible that the selection pressure for species discrimination traits in guenons arises partially or entirely from other functions where behavioral coordination or avoidance between species is advantageous, such as in foraging decisions [20,23]. Careful field observations would be needed to distinguish between such possibilities.

Our occlude-reclassify approach is a novel method for identifying the distribution of information in complex signals and can be used for any question that can be conceptualized as a discrimination problem and analyzed using machine classification. This method therefore has broad utility within sensory ecology and could help to better understand the link between form and function in the complex signals that are common in many animal groups. The objectivity of the approach is important, as it allows researchers to intelligently target specific signal components for further analysis without reference to their own perceptions of their salience. This is particularly important when studying species with sensory and perceptual systems very different from our own [50,51]. Where possible, combining this approach with a biologically realistic classification scheme, such as classification within a perceptual face space based on eigenface scores [36] as used here, increases the biological validity of results.

Our research broadens our understanding of how morphology and social decision-making can interact to structure interactions between species living in sympatry. In guenons, facial features like white nose spots are highly salient, attention-grabbing, and distinctive, and our combined results demonstrate the importance of these traits in species discrimination. Guenon behavioral repertoires, such as nose-to-nose touching observed in wild putty nosed monkeys (SW, personal observation) and red-tailed monkeys [52], further reflect the importance and biological relevance of these traits. Primates preferentially attend to facial information [53,54], making face patterns particularly suited to influencing behavior and decision-making in con- and heterospecifics. The evolution of signals facilitating species discrimination may be a major driver of biological diversity, and our work linking mating signal form and function in a recent and diverse primate radiation highlights how such evolutionary processes can be important in generating animal phenotypes.

## Supporting information

Supplementary Results

## Acknowledgements

We thank Allegra Depasquale and Laura Newman for assistance coding looking time videos, and Kathryn Yee for assistance with stimuli preparation. Special thanks the directors and staff of CERCOPAN sanctuary, particularly Claire Coulson and Isabelle Theyse, for access to the facility and support during data collection.

## Data accessibility

The guenon face image database, experimental looking time data, and all project code will be uploaded to Dryad. During review, all files are available at: http://www.dropbox.com/sh/1yf97kbhggzi5a8/AAAJ0dQMEqeFOMf9aHzGBMHNa?dl=0

