## Supplementary Methods for "The structure of species discrimination signals across a primate radiation"

### Occlude-reclassify procedure

In this study, we used an occlude-reclassify procedure to identify face regions critical for correct machine classification of guenon faces by species. The general approach of this procedure is straightforward: different signal components (i.e. face regions in this study) are systematically occluded and the image reclassified; essential signal components are those that cause mis-classification when occluded. However, several details must be considered when implementing this procedure using images. We overview some of these here.

*Occluder color*

The occluder that masks each image region must generally be set to a color, and there are multiple options for this. The goal of the procedure is to negate the given region, and the best method of doing so will differ across circumstances. In some cases, it may be possible to model the occluded region as missing data (e.g. as NaN in MATLAB), however this can introduce computational problems (e.g. the PCA-based method used here requires complete data) or complicate interpretations (e.g. there are no missing data in natural scenes, making this less biologically realistic). Here, we employed two different occluder colors: (1) the average grey of all guenon faces (i.e. the color used for the background of all images), and (2) the mean face color (calculated without including the background) of the species in question. Average grey occluders have been used previously [1], are convenient to implement, and generate results that are directly comparable across groups. However, using the same color occluder for all groups potentially biases classification performance as classification should be better for species that have more average color faces than species with very light or dark faces. Furthermore, in this project we compared computational results derived using the occlude-reclassify procedure to results from experiments in which facial stimuli were presented to guenon subjects. In those experiments some facial stimuli were modified, with certain face regions (either nose spots or eyebrow patches) altered to better match the rest of the face. A species-specific occluder color is more in line with these experimental manipulations and therefore present these results in the manuscript. In setting the color of the occluder to the species-specific mean face color we introduce a different potential bias, effectively increasing the likelihood of detecting face regions that are salient against the general face color of the species. However, these regions will also be more salient to receivers, especially since our analyses are conducted in guenon color space. Given that our goal is to identify face regions most likely to contribute to species discrimination, this bias seems reasonable. For completeness, we have also included results generated using a standard average grey occluder (see supplementary results).

*Occluder size*

The size of the occluder must also be specified. This choice reflects a trade-off: for very small occluders, misclassification will be very rare and essential signal components will not be identified; for very large occluders, misclassification will be very common and the identified signal components will converge on the entire area of the signal. We chose an occluder size that balanced these two extremes, being large enough to detect face regions of interest, but small enough that the regions identified were restricted to only some regions of the face and therefore useful for identifying critical facial features.

*Occluder shape*

Occluders can also vary in shape. In our analyses we used a square occluder, which will often be the most reasonable choice. Other shapes may sometimes be more useful, however, depending on the characteristics of the signal of interest. For instance, long rectangular occluders may be useful for analyzing stripes, or more complex patterned occluders (e.g. multiple spots) for analyzing signals with repeated pattern elements.

A potential alternative use of the occlude-reclassify procedure is to test multiple occluders of different shapes to determine which causes the most disruption of correct classification and therefore represents the most biologically meaningful pattern.

*Signal symmetry*

Some signals, such as the faces analyzed here, are symmetrical. Sliding an occluder across an entire image that depicts a symmetrical signal may be problematic because mirrored features will never be fully occluded. For instance, if cheek tufts in one species and a nose spot in another species are equally critical for correct classification, cheek tufts – which exist in pairs on opposite sides of the face – may nonetheless be less likely to be identified using the occlude-reclassify procedure because when one is occluded the other is still visible to the classifier. By comparison, a single nose spot in the middle of the face could be occluded in its entirety, completely masking this information and potentially making this feature more likely to be identified as critical using this scheme. We address this by classifying hemi-faces, in which the signal is split along the line of approximate symmetry. In our implementation of the occlude-reclassify procedure the occluder location is recorded based on the central pixel, so there is a buffer along the edges of the images (equal to one half of the occluder length) in which critical signal regions are not detected. The edges of the full images are simply grey background, so this is not problematic, however when generating hemi-face images a buffer region must be added alongside the midline of the face to allow the occluder to reach the center of the face. In order to give midline regions of the face equal importance in the classification we used the first 15 columns of the other half of the face as a buffer. Prior to analysis we arbitrarily chose to present results based on the left hemi-face (from the perspective of the viewer; the right side of the animal's face). These are similar to results based on the left hemi-face, although not identical (see supplementary results).

**Looking time experiments**

*Experimental guenon population*

Looking time experiments were conducted at CERCOPAN sanctuary in Calabar, Nigeria, and included 18 adult putty nosed monkeys (*C. nictitans*; n males = 6, n females = 12; mean (range) age = 13 (7-22) years) and 16 adult mona monkeys (*C. mona*; n males = 10, n females = 6; mean (range) age = 8.8 (3-24) years). All subjects received appropriate primate diets, environmental enrichment, and veterinary care, and were socially housed with other guenons in groups of two to five (mean = 4). All subjects were within visual range of heterospecific guenon species, which included *C. erythrotis*, *C. mona*, *C. nictitians*, *C. preussi*, *C. sclateri*, *Ch. tantalus*, and *E. patas*, although the exact heterospecific species visible varied by group. Primate species housed at CERCOPAN are all endemic to Nigeria. Each subject was therefore familiar with both conspecific and sympatric heterospecific guenon faces.

Each species was divided into four experimental groups, which served as experimental replicates. Many experimental groups were also complete social groups, however in some cases social groups were combined to create experimental groups (e.g. two social groups containing only two mona monkeys were combined to yield an experimental group containing four subjects). Each experimental group received the same experimental treatments; i.e. they viewed the same stimulus image pairs in the same order. Putty nosed monkey experimental groups contained either four or five subjects (mean = 4.5), and all mona monkey experimental groups contained four subjects.

*Stimulus image preparation*

We obtained stimulus images from our image database of guenon faces, which were prepared in a similar way to those used for computational analyses. We cropped images to display the head and shoulders of the subject; applied color constancy using a combination of the gray world assumption and maximum point of reflectance, as described previously [2]; standardized blur across images to a blur metric [3] of 0.3; and transformed images such that the outer corners of the eyes were level and 150 pixels apart. Standardized naturalistic backgrounds are preferable in looking time tasks [4], therefore we segmented the subject from the background and replaced the latter with a blurred image depicting natural guenon forest habitat (taken in Gashaka Gumti National Park, Nigeria). We did not use a standardized reference frame (e.g. using the eyes as landmarks to standardize the positioning of the face) across all stimuli because this yielded images that were strangely framed due to the variability in facial characteristics and positioning (e.g. an excessive amount of 'background' space on some sides of the face). Instead we positioned the subject on the background image such that the framing appeared natural. This processing pipeline yielded a standardized set of stimulus images presented at approximately life-size. Stimulus image manipulations were conducted using the Image Processing Toolbox in MATLAB [5] and the GNU Image Manipulation Program [6].

We matched each stimulus image pair for sex, with each subject viewing images of either all males or all females to facilitate comparisons of subjects across trials. We were unable to standardize eye gaze across all stimulus images, and so accounted for this factor statistically by incorporating stimulus eye contact as a variable in our analysis. All stimuli depicted mature adults without any type of aberrant facial appearance (e.g. scars, abnormal facial characteristics, hair loss, etc.). Stimulus images presented to mona monkeys were all from individuals outside CERCOPAN and were therefore all unfamiliar. CERCOPAN is the only source of putty nosed monkey face images in our database, therefore for their trials it was necessary to use images collected on site as stimuli. We minimized the familiarity between subjects and animals represented in stimulus images as much as possible; in some cases it was necessary to present a stimulus image of an animal in visible range, however this was always at a distance and subjects never saw stimuli of animals in their own or adjacent cages. To account for any impact of individual familiarity we included this as a binary variable in statistical analyses of putty nosed monkey eye gaze (not within visible range vs. within visible range).

Our stimulus image set consisted of four images (2 male, 2 female) of each heterospecific species: *C. ascanius*, *C. diana*, and *C. wolfi*; six images (3 male, 3 female) of mona monkeys; and twelve images (6 male, 6 female) of putty nosed monkeys. Mona monkey subjects were presented with all relevant stimulus images of one sex (each experimental group saw either male or female images). Putty nosed monkey subjects were presented with all relevant heterospecific stimulus images of one sex, but each experimental group was presented with unique putty nosed monkey stimulus images (sex-matched to heterospecifics) to reduce familiarity effects as much as possible.

*Experimental apparatus*

We designed an experimental apparatus (supplementary figure 1) that required guenons to view stimuli through a viewing window, effectively forcing them to remain relatively stationary while participating in experiments. Our experimental apparatus consisted of an opaque box (58 x 48 x 35 cm) housing all experimental equipment, with a small (14 x 6 cm) subject viewing window on one side. Inside the box we positioned a widescreen laptop (Acer ES1-711-P1UV) opposite the viewing window and a video camera (Canon Vixia HF R40 HD) centered immediately above the laptop screen to record the subjects' responses. The laptop LCD screen was color characterized weekly using ColorMunki Design [7], with the resulting color profile used to ensure the accurate presentation of colors in all stimulus images. To help interest subjects in the experimental apparatus, the external side of the apparatus that contained the viewing window was painted with a colorful pattern using watercolors. The pattern was varied across trials to facilitate sustained interest throughout the duration of experiments. Subjects could not see this colorful portion of the apparatus when looking through the viewing window, but we nonetheless included the external pattern used for the trial in statistical analyses to account for any impact this may have had on looking biases. The video camera records both video and sound, allowing researchers to dictate the names of subjects who look inside the apparatus to the camera. To minimize distractions, we typically said subject names aloud after subjects had moved away from the apparatus. Most subjects were inherently interested in the experimental apparatus, however in some cases food rewards were used to encourage participation ($n = 5$ trials involving $n = 3$

subjects). Food rewards included peanuts or diluted juice presented in a water bottle, selected based on subject preferences. The presence of food is unlikely to influence species discrimination capabilities or visual biases for faces.
