## Supplementary Results for "The structure of species discrimination signals across a primate radiation"

For completeness, we ran our occlude-reclassify procedure using two occluder colors and both left and right hemifaces. For an explanation and justification of these choices see supplementary methods. We present results generated using species-specific mean face colored occluders and left hemifaces in the results section; here, we present and discuss the remaining results.

We used two occluder colors to satisfy different conditions: a species-specific mean face color occluder is consistent with the goal of the study to identify how discrimination is supported by signal traits in different species, however using the same grey occluder across species facilitates comparisons across species. Results based on the species-specific mean color (left hemi-faces: figure 3; right hemi-faces: supplementary figure 2) and the average grey (left hemi-faces: supplementary figure 3; right hemi-faces: supplementary figure 4) are very similar for some species. For instance, the results for *C. ascanius* and *C. lhoesti* are virtually identical using the two methods; this is unsurprising because the mean face color for each of these species is very similar to the overall average face color used for the grey occluder. However, for species with generally darker or lighter faces, results differ between the two methods. For instance, *M. talapoin* and *C. nictitans* are among the lightest and darkest faced guenon species, respectively, consequently results differ depending on occlude color. Again, this is unsurprising because the occluder itself is salient against these faces, rather than muting a face region as in other species. In these cases the grey occluder is less realistic, thus our choice to present the species average face color results in the main section of this manuscript. It is worth noting that our experimental results documenting putty nose monkey (*C. nictitans*) visual biases are much more strongly paralleled by occlude-reclassify results based on a species-specific mean face color occluder. This is particularly relevant because the modifications made to putty nosed monkey stimulus images used in experiments involved darkening the white nose spot to match the rest of the face, a procedure that corresponds to using a darker occluder.

We also ran occlude-reclassify analyses using right and left hemi-faces separately, to account for any effects of bilateral symmetry in guenon faces (see supplemental methods). Across species there are broad similarities in the parts of the face identified as critical for correct species classification using the left (species-mean occluder: figure 3; grey occluder: supplementary figure 3) and right hemi-faces (species-mean occluder: supplementary figure 2; grey occluder: supplementary figure 4). In some cases a feature is identified as having greater relative importance (i.e. a higher rate of misclassification) on one side compared to the other (e.g. the eyebrow region of *C. wolfi*), and in others a feature is only identified on one side (e.g. the cheek tufts of *Ch. sabaeus*). Differences are likely due to asymmetries in posture introduced because of the need to photograph unrestrained and mobile animals. More pronounced differences are found in some species with low sample sizes, reinforcing that results for these species should be interpreted with caution. Despite some variation in results from each side of the face, the general patterns observed in our occlude-reclassify results are similar and our conclusions remain unchanged. Importantly, results from both putty nosed monkeys (*C. nictitans*) and mona monkeys (*C. mona*) are very similar for the two hemi-faces, with our classifier relying exclusively on the nose spot (albeit with a stronger effect observed in the right hemi-face) and across broad face regions, respectively. Therefore, variation between results generated based on different hemi-faces does not influence our interpretations of the parallels between computational and experimental results in these two species.

### Supplementary Figures & Tables

**Supplementary Figure 1**

The experimental apparatus. (a) The apparatus was painted to draw interest and placed immediately outside subject enclosures. (b) A widescreen laptop and video camera were housed inside the apparatus to display stimulus images and record subject responses. (c) View of subject participating in looking time trial as recorded by the internal video camera.

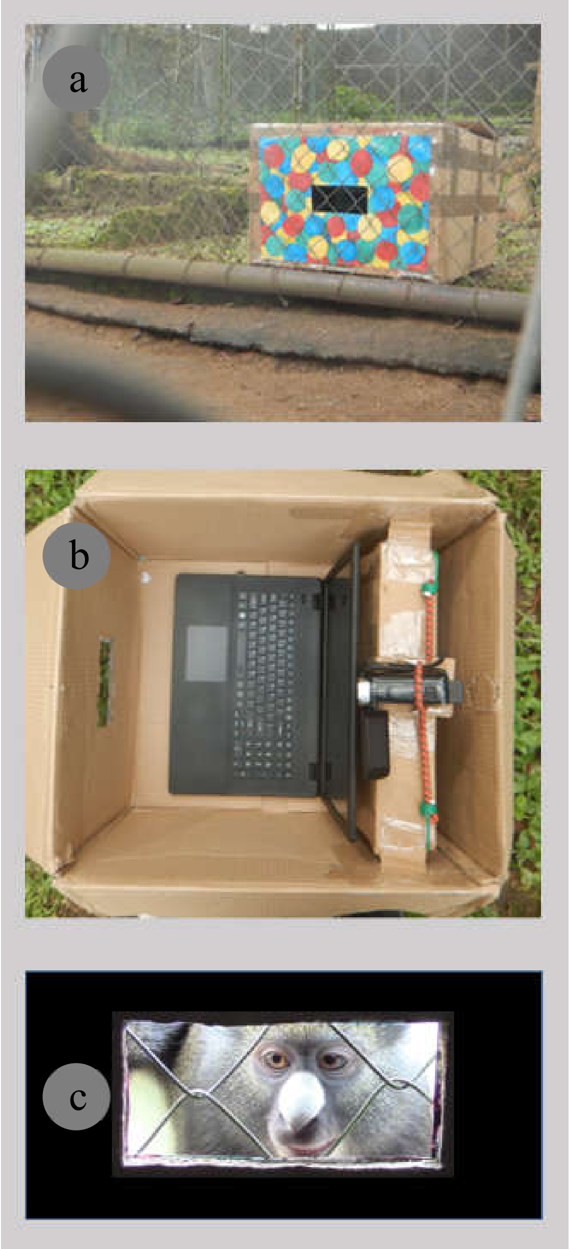

**Supplementary Figure 2**

Likelihood of correct classification based on occlusion of different face regions with the occluder set to the species-specific mean face color and run on the right hemi-face (see Supplementary Methods). Species average faces are displayed on the left and heatmaps identifying critical face regions on the right. Sample size is reported as n = number of individuals (number of total images).

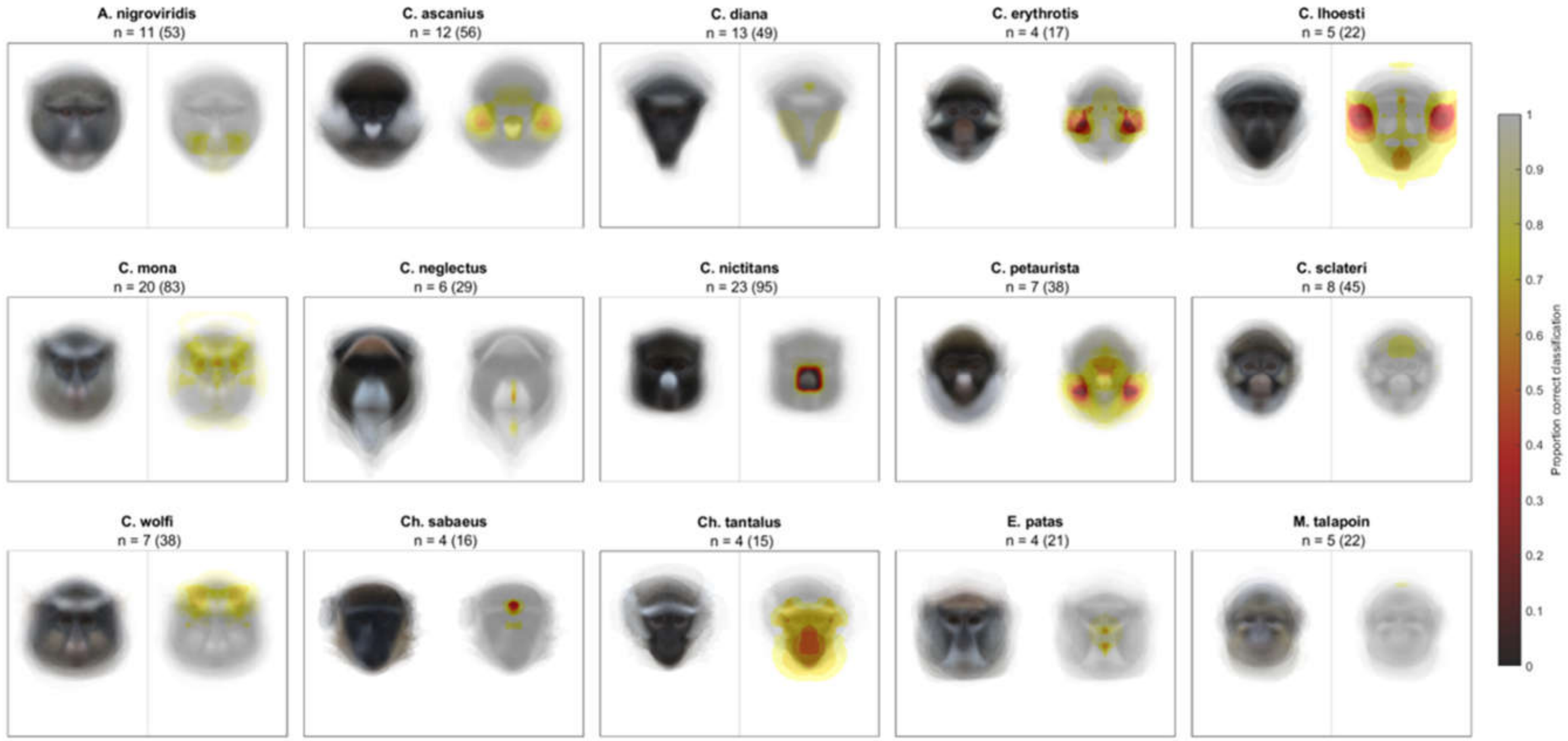

**Supplementary Figure 3**

Likelihood of correct classification based on occlusion of different face regions using an average grey occluder on left hemi-faces (see Supplementary Methods). Species average faces are displayed on the left and heatmaps identifying critical face regions on the right. Sample size is reported as n = number of individuals (number of total images).

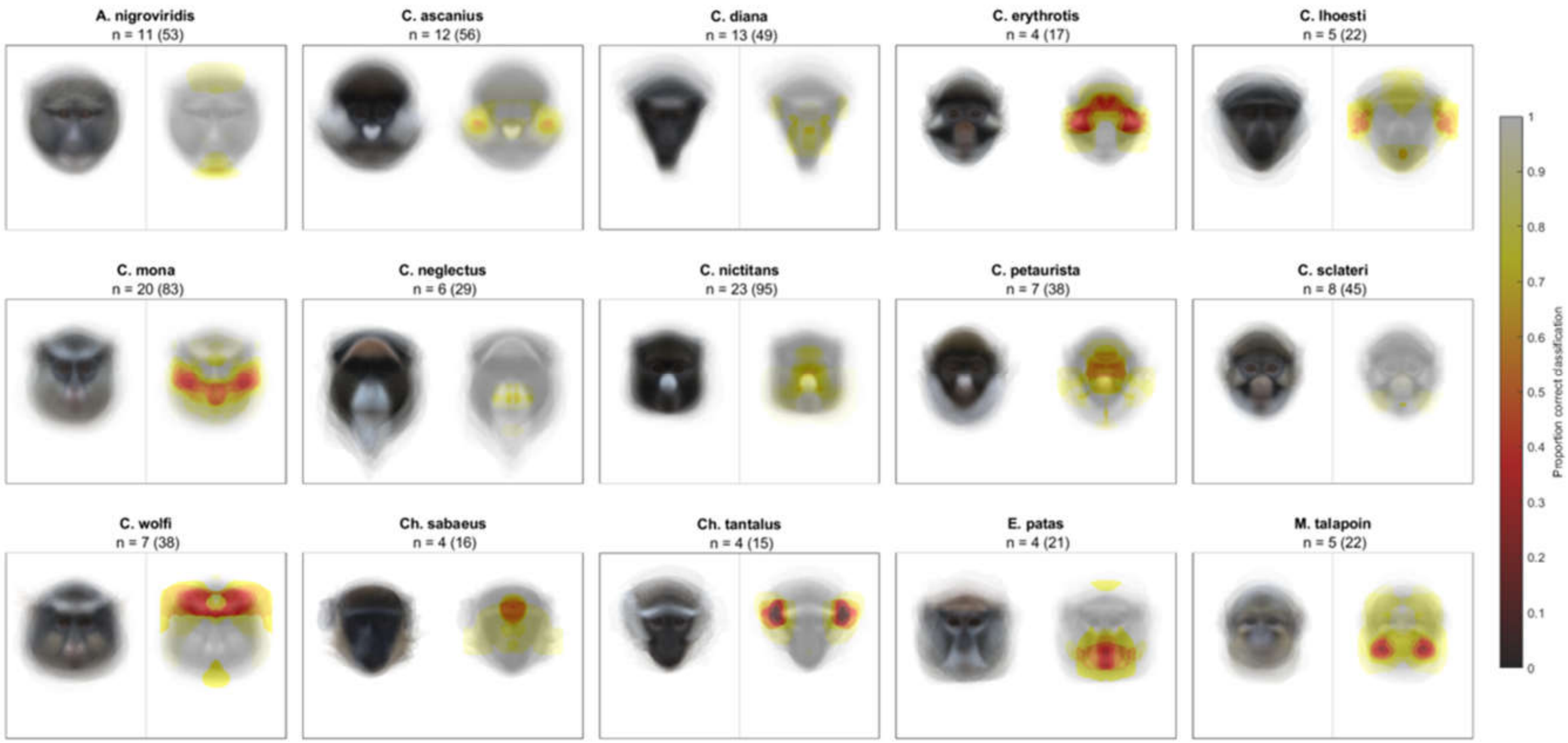

**Supplementary Figure 4**

Likelihood of correct classification based on occlusion of different face regions using an average grey occluder on right hemi-faces (see Supplementary Methods). Species average faces are displayed on the left and heatmaps identifying critical face regions on the right. Sample size is reported as n = number of individuals (number of total images).

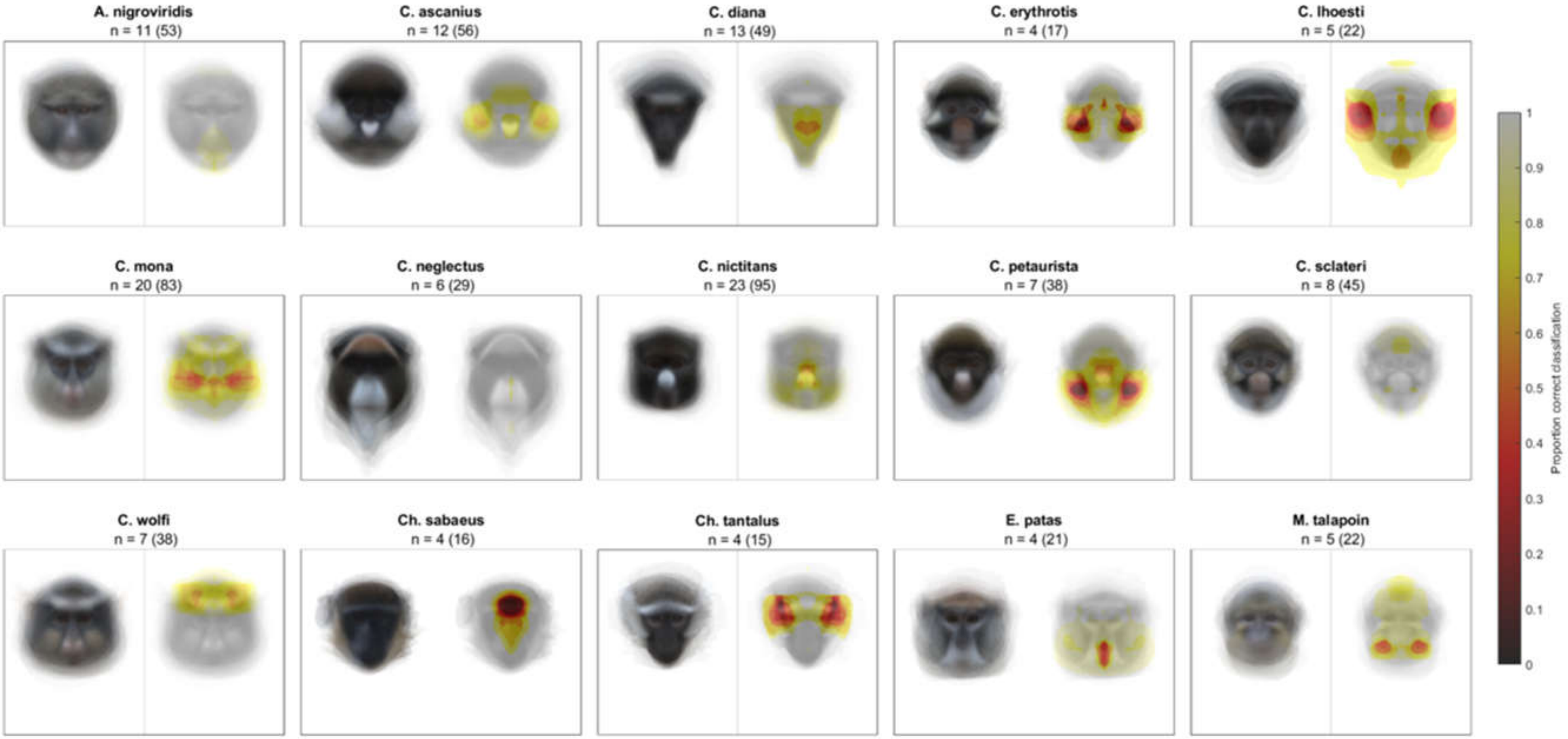

**Supplementary Table 1**

Results from model comparisons for putty nosed monkeys. The final model is the best fit to the data and was used to make factor-level comparisons.

| Model | Fixed effects | Comparison | Likelihood ratio test | |
|---|---|---|---|---|
| M1 | None | n/a | n/a | |
| M2 | Species (con- v. heterospecific) | M1 | Chisq = 3.051, df = 1, p = 0.081 | · |
| M3* | Trait (shared v. not shared) | M1 | Chisq = 10.473, df = 1, p = 0.001 | ** |
| M4 | Subject sex (male v. female) | M1 | Chisq = 0, df = 1, p = 1 | |
| M5 | Subject age (numerical) | M1 | Chisq = 0, df = 1, p = 1 | |
| M6 | Subject origin (captive v. wild) | M1 | Chisq = 0, df = 1, p = 1 | |
| M7* | Presentation side (right v. left) | M1 | Chisq = 17.697, df = 1, p < 0.001 | *** |
| M8 | Eye contact (yes v. no) | M1 | Chisq = 0.185, df = 1, p = 0.667 | |
| M9 | Familiarity (ordinal: not present < present but not visible < visible) | M1 | Chisq = 3.878, df = 2, p = 0.144 | |
| M10 | Stimulus sex (male v. female) | M1 | Chisq = 0, df = 1, p = 1 | |
| M11 | Trial order (categorical: first, second, third) | M1 | Chisq = 0, df = 2, p = 1 | |
| M12 | Apparatus pattern (4 categories) | M1 | Chisq = 0, df = 3, p = 1 | |
| M13 | ICC profile (5 categories) | M1 | Chisq = , df = 1, p = | |
| M14* | Species + trait + presentation side | M1 | Chisq = 31.933, df = 3, p < 0.001 | *** |
| M15 | Trait + presentation side[1] | M14 | Chisq = 3.055, df = 1, p = 0.081 | · |
| M16 | Species + presentation side[1] | M14* | Chisq = 11.511, df = 1, p < 0.001 | *** |
| M17 | Species + trait[1] | M14* | Chisq = 18.065, df = 1, p < 0.001 | *** |
| Final* | Trait + presentation side | M1 | Chisq = 28.878, df = 1, p < 0.00 | *** |

*Denotes which of the two models was a better fit to the data when the difference was statistically significant.
[1]Model comparisons test the significance of the factor that was removed from the expanded model (e.g. comparing M15 to M14 tests the significance of species).

**Supplementary Table 2**

Results from model comparisons for mona monkeys. The final model is the best fit to the data and was used to make factor-level comparisons.

| Model | Fixed effects | Comparison | Likelihood ratio test | |
|---|---|---|---|---|
| M1 | None | n/a | n/a | |
| M2[*] | Species (con- v. heterospecific) | M1 | Chisq = 179.640, df = 1, p < 0.001 | *** |
| M3[*] | Trait (shared v. not shared) | M1 | Chisq = 31.621, df = 1, p < 0.001 | *** |
| M4 | Subject sex (male v. female) | M1 | Chisq = 0, df = 1, p = 1 | |
| M5 | Subject age (numerical) | M1 | Chisq = 0, df = 1, p = 1 | |
| M6 | Subject origin (captive v. wild) | M1 | Chisq = 0, df = 1, p = 1 | |
| M7 | Presentation side (right v. left) | M1 | Chisq = 0.985, df = 1, p = 0.321 | |
| M8 | Eye contact (yes v. no) | M1 | Chisq = 0.761, df = 1, p = 0.383 | |
| M9 | Stimulus sex (male v. female) | M1 | Chisq = 0, df = 1, p = 1 | |
| M10 | Trial order (categorical: first, second, third) | M1 | Chisq = 0, df = 2, p = 1 | |
| M11 | Apparatus pattern (4 categories) | M1 | Chisq = 0, df = 3, p = 1 | |
| M12 | ICC profile (4 categories) | M1 | Chisq = 0, df = 3, p = 1 | |
| M13[*] | Species + trait | M1 | Chisq = 209.100, df = 2, p < 0.001 | *** |
| M14 | Trait[1] | M13[*] | Chisq =177.480, df = 1, p < 0.001 | *** |
| M15 | Species[1] | M13[*] | Chisq = 29.462, df = 1, p < 0.001 | *** |
| M16[*] | Species*trait + species + trait | M13 | Chisq = 8.242, df = 1, p = 0.004 | ** |
| Final[*] | Species*trait + species + trait | M1 | Chisq = 217.340, df = 3, p < 0.001 | *** |

[*]Denotes which of the two models was a better fit to the data when the difference was statistically significant.
[1]Model comparisons test the significance of the factor that was removed from the expanded model (e.g. comparing M14 to M13 tests the significance of species).
